# Characterizing Industrial Pond Ecology Timeline in DISCOVR Cultivation Trials for Early Detection of Pond Crashes

**DOI:** 10.64898/2026.03.31.715673

**Authors:** Elise K. Wilbourn, Deanna Curtis, Hemanth Kolla, Prashant Madan Mohan Rai, Pamela Lane, John McGowen, Todd W. Lane, Kunal Poorey

**Author notes:** Corresponding author: Kunal Poorey, 7011 East Ave, Livermore CA 94550.

## Abstract

For sustainable algal biomass cultivation, we need substantial improvement in annualized productivity by reducing the frequency of crop failure and improved growth in open raceway pond systems. In this study, high-performing strains were identified and optimized for biomass productivity. We utilized next-generation sequencing methods to quantify the ecological features of open raceway systems cultivated at in Arizona. We utilized data from several months of cultivation runs to construct a rich time-series of the ecology dynamics using amplicon sequencing and used custom anomaly detection, “PondSentry”, for the early prediction of pond crashes. PondSentry uses tensor decomposition of higher-order joint moments to detect incipient anomalies in multivariate data and displays significant improvements from standard knowledge-based anomaly detection methods. The PondSentry strategy identifies signs of deteriorating pond health at an average of three days before an actual crash event, with rank order of the ecological features plausible for crop failures driven by organisms such as *Amoeboaphelidium occidentale* FD01. These findings are independently confirmed with PCR and microscopy studies at an Arizona cultivation site. PondSentry’s time-series-based anomaly detection of crashes provides a suitable monitoring strategy for eukaryotic crash agents in unialgal culture. The early warnings can be used to time interventions or harvests to prevent biomass loss. The PondSentry strategy strengthens the role of data science and data-driven methods in algal cultivation and can increase the feasibility of algal-biomass based products.

## Introduction

Alongside other biomass production methods, algae-based biomass grown in multiacre outdoor ponds shows promise on the path to United States energy independence [1-3]. In addition to biofuels, algal biomass can be utilized for pharmaceuticals, food, nutraceuticals, wastewater management, animal feed, bioproducts, and many other applications [4-8]. For sustainable algal biomass cultivation, we need substantial improvement in annualized productivity by reducing the frequency of crop failure and improving growth on open raceway systems [9-12]. In industrial algae production, an estimated 10-30% of biomass loss is due to biological contamination (from infections, pathogens, and grazers)[13]. Effective monitoring and identification of these deleterious agents will help identify crop protection strategies and decisions on pond operating strategies for biomass loss prevention.

Current monitoring strategies are often time- and resource-intensive, requiring direct human monitoring via microscopic and molecular methods. This both adds to the final capital expenditure of any algae-based products and leaves room for human error. In addition, these methods often detect algae population crashes driven by contaminants hours to a few days prior to the crash, limiting response time. Finally, while pathogens including fungi and grazers are visible via microscopy, determination of causative agents is often a lengthy process, limiting the response to broad-spectrum interventions such as complete crop harvest rather than targeted methods tailored to the specific pest [14, 15].

Here we report results from several months of outdoor, unialgal raceway cultivation as interpreted with the data-science toolkit “PondSentry”, which can be used to predict early indicators of algal pond crashes and times of low productivity. PondSentry uses tensor decomposition of higher-order joint moments to detect incipient anomalies in multivariate data and displays significant improvements from standard knowledge-based anomaly detection methods. This method can detect pond crashes up to four days before the lowest productivity associated with pond crashes, and can give rank orders of causative agents of the pond crash. This was shown here with the detection of fungal pests including *Amoeboaphelidium occidentale* FD01[16], with confirmation of the crash validated with PCR and microscopy studies at the Arizona Center for Algal Technology (AzCATI). PondSentry uses a time-series-based anomaly detection strategy to flag eukaryotic crash agents in unialgal raceway cultures. Early warnings allow for longer reaction times to either implement other crash interventions or harvest to prevent biomass loss. The PondSentry strategy strengthens the role of data science and data-driven methods in algal cultivation and shows great promise for future crop monitoring systems.

## Methods and Materials

### Algal Cultivation

Algae were grown in open raceway ponds at the Arizona Center for Algae Technology and Innovation (AzCATI) at the Arizona State University Polytechnic Campus in Mesa, AZ. Pond cultivation and sequencing methods have been previously published [17] and so will only be briefly described here. Two strains of green algae were grown during this study period, *Tetradesmus obliquus* UTEX393 and *Monoraphidium minutum* 26B-AM[18]. Samples were collected from ponds between Nov 4^th^ and Dec. 30^th^, 2019. Grab samples were taken from each pond in 15 mL sterile polycarbonate tubes (Falcon) and stored at -20 °C with 2% RNALater (Sigma Aldrich) prior to analysis. Samples were shipped frozen from AzCATI to Sandia National Laboratories in Livermore, CA for genetic analysis with next generation sequencing methods. Sequencing occurred no more than one year post-collection. DNA was extracted following manufacturer instructions with the Zymo Quick-DNA Fungal/Bacterial 96 Kit.

### Library preparation and sequencing

The 18S V4 sequencing libraries were prepared via nested PCR by first amplifying the V4 hypervariable regions of the eukaryotic 18S ribosomal DNA using the primer pair TAReuk454_F and TAReukREV3 [19]. The first PCR amplification reaction contained 22.5 µL of AccuPrime Pfx Supermix (ThermoFisher Scientific), 0.2 µM final concentration of each of the TAReuk454_F and TAReukREV3 primers, and 12.5 ng of genomic DNA in a final reaction volume of 26 µL. The cycling conditions were an initial denaturation at 95 °C for 5 min; 10 cycles of 95 °C for 80 s, 57 °C for 90 s, and 65 °C for 1 min; 25 cycles of 95 °C for 80 s, 48 °C for 90 s, and 65 °C for 1 min; and a final 5-min extension at 65°C. PCR products were cleaned and concentrated to 20 µL using a Zymo DNA Clean & Concentrator kit.

The second round of PCR was performed to add Illumina-specific index sequences. The second PCR amplification was carried out on 2.5 µL of V4 amplified DNA product from the first PCR reaction along with 12.5 µL of Failsafe Buffer E (VWR, PA, USA), 0.25 µL of Failsafe Enzyme, 4.75 µL nuclease free water, 2.5 µL of the octonucleotide Illumina primer 1, and 2.5 µL of the octonucleotide Illumina primer 2 [Gloor et al, 2010] in a 25 µL total reaction volume. The amplification conditions were taken from the 16S Metagenomic Sequencing Library preparation: denaturation at 95 °C for 3 min, 8 cycles of 95 °C for 30 s, 57 °C for 30 s, and 72 °C for 30 s; and a final 5-min extension at 72 °C. PCR products were cleaned and concentrated to 20 µL using a Zymogen DNA Clean & Concentrator kit, as above.

Illumina-barcoded DNA was quantified using the Quant-iT dsDNA Assay Kit, as above, and 50 ng of 96 samples were combined. The V4 library average DNA size was determined to be 578 bp with a 2100 Agilent Bioanalyzer (Agilent, Santa Clara, CA, USA). Sequencing was performed with a V3 300-cycle paired end kit (Illumina, San Diego, CA, USA) on an Illumina MiSeq. The denatured library was diluted to 6 pM in HT1 buffer (Illumina) and mixed with 10% PhiX (6 pM).

### Bioinformatics Analysis of Amplicon Sequencing -part 2

We applied bioinformatics pipelines MAGPie and RapTOR for estimating 18S taxonomic abundances as OTUs for ∼100 samples. Reads were classified against the Silva and Greengenes library. This library targets primarily eukaryotic species.

## Results and Discussion

### Bioinformatics Analysis of Amplicon Sequencing

We first estimated eukaryotic taxonomic abundances for 110 samples. We use raw 18S amplicon sequencing data and output taxonomic abundance tables. A summary of taxonomy classification in the UTEX393 cultivation run from October 30 to December 31, 2019 is shown in Figures 1 and 2. As we are only interested in the contamination in the SOT run, we removed the reads coming from the seeded algae strains and only kept reads from possible contaminants. Figure 1A shows the broad characterization of the contaminant reads at the kingdom level. The pattern is best represented as a heatmap to highlight the rise and fall of contaminants in the ponds over the cultivation time course. We chose three ponds (numbered 23, 24, and 26) to best capture the timeline of the biological anomaly (crash) event. Major categories of the contaminants in these ponds include *Fungi, Metazoa, Aphelida*, and other phototrophic contaminants.

**Figure 1:**
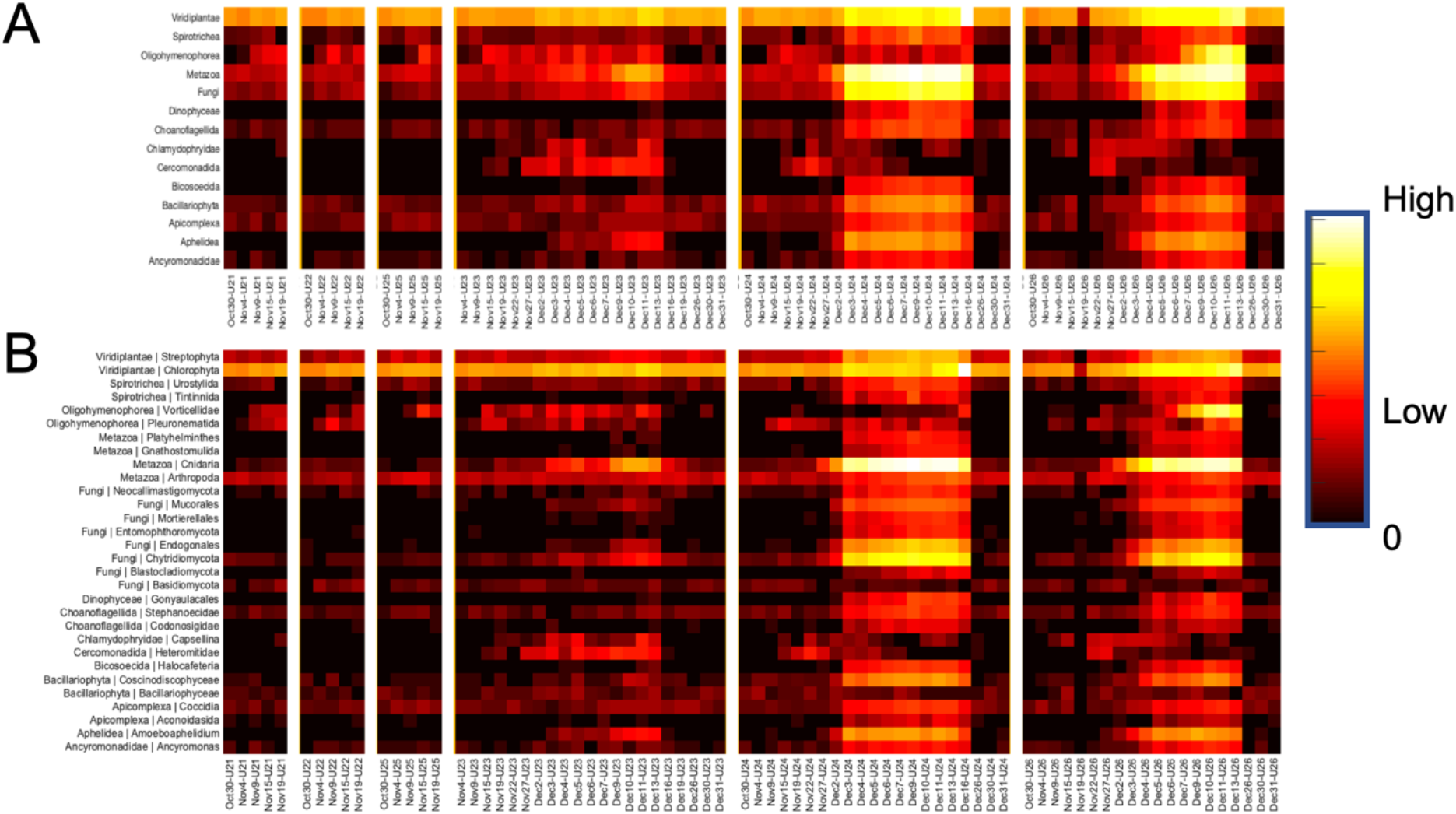
Broad classification of contaminant reads from the SOT cultivation runs. A: shows a heatmap of log-scale abundances of reads at kingdom level, and B: shows a heatmap of log-scale abundances of reads at division level.

**Figure 2:**
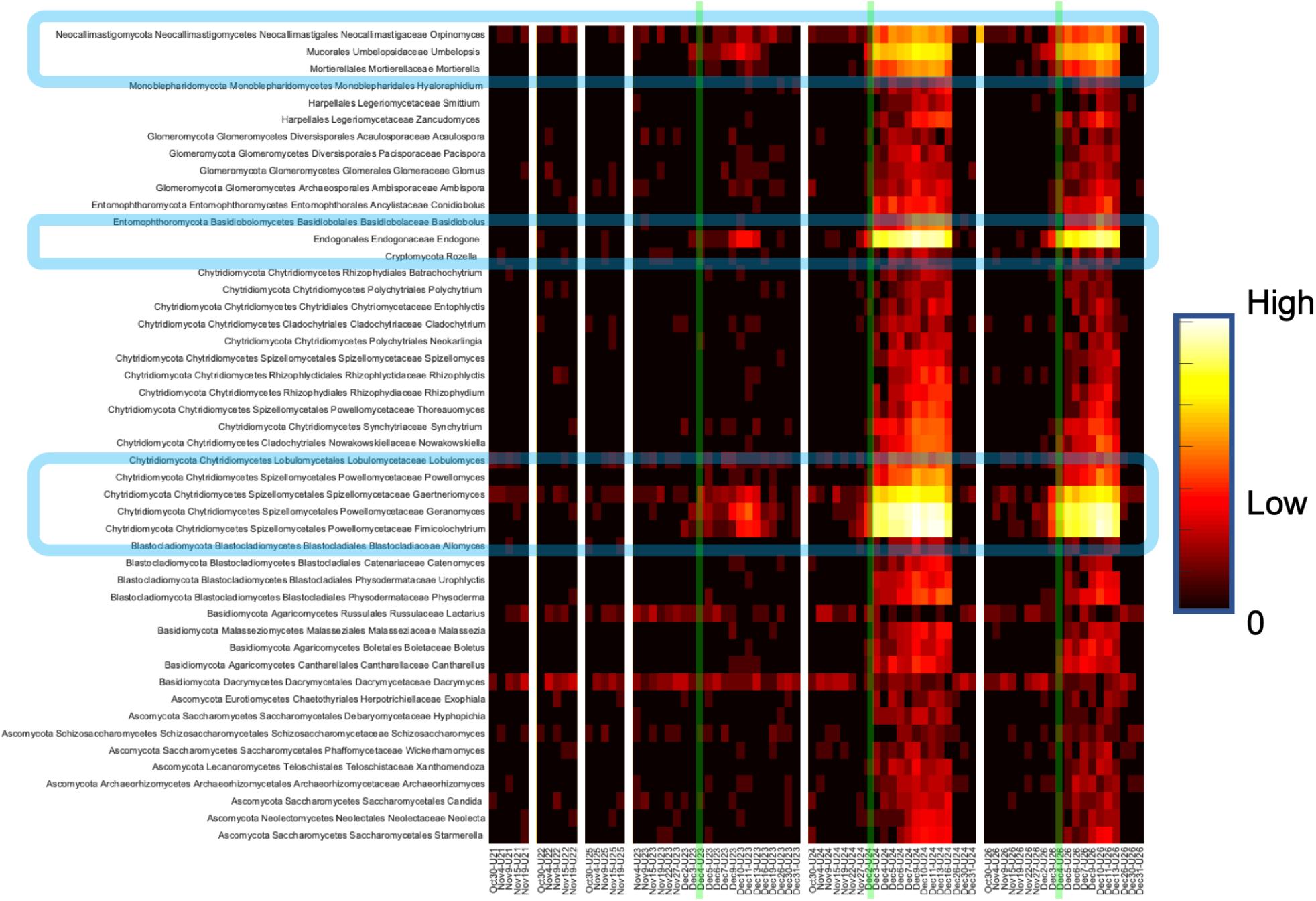
Heatmap of log-scale amplicon sequencing abundance classified to the various fungal genera. The dominance of Chytrid fungus in these taxonomy can be observed from Dec 5th onwards in Pond 23, 24 and 26.

Figure 1B shows a time series of these contaminants. Prominent groups include the fungi *Chytridiomycota*, the fungus-like *Aphelidea*, and *Cnidaria* (which can be traced to “*hydra,”* a freshwater organism), along with other phototrophic contaminants. Figure 2 shows the classification of the fungal genera present in the samples.

As with other crashed cultivation ponds, fungi were present as contaminants in this system. The analysis showed 48 significant genera of fungi were present in the cultures, which are presented in Figure 2. This figure displays a log scale representation of fungal abundances present in the samples, with blue boxes highlighting key community members. For all three ponds, the abundance of the contaminant increases around Dec 4th to Dec 5th. The rise is more dramatic in Ponds 24 and 26 than the Pond 23. The majority of the reads (at an average >50% of the reads) in the fungal genera are contributed from various Chytrid fungi (as shown in Figure 2 as a Heatmap). However, it is not clear from genetic data alone whether the fungi present were responsible for the pond crash. While members of *Chytridiomycota* have been implicated in crashes in other strains of algae, they are also ubiquitous in environmental systems, evidently without causing algal population crashes [20, 21].

We also explored how the eukaryotic diversity changed in these ponds over the cultivation timeline. Figure 3 shows the Shannon diversity index plotted through the time course of cultivation for the three ponds, all of which reported crashes. We observed an increase in the diversity index at the peak of the pond crash, which was expected. Figure 4 shows Krona charts for the taxonomic abundance and distribution for selected samples for in-depth analysis and pond ecology exploration. These plots also demonstrate patterns in diversity over time. We show that as of November 19^th^, the majority of the reads were coming from the UTEX393 strain. By December 2^nd^, we detect some fungal contamination and grazers. By December 10^th^, the number of reads from fungus, grazers, and other eukaryotic contaminants increases at the peak of the crash. The final plots shown are from December 26^th^ and show a signature of a clean pond due to a reset from a clean algae seed train. In addition to overall patterns in diversity, we show that there were differences in the overall diversity of the ponds compared to each other. Pond 26 displayed significantly lower baseline diversity than the other two ponds.

**Figure 3:**
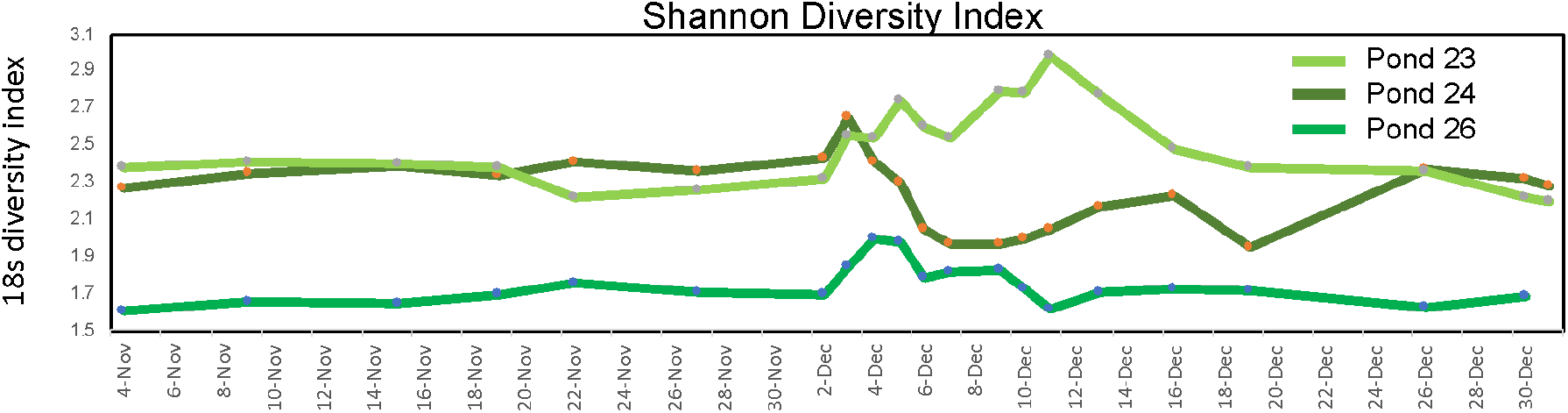
We showcase the dynamics of the eukaryotic diversity measured through Shannons Diversity Index for the timeline of the cultivation run. There is a noticeable rise in the eukaryotic diversity between Dec 3rd and Dec 12th. This pattern is expected as eukaryotic diversity correlates with crashed samples.

**Figure 4:**
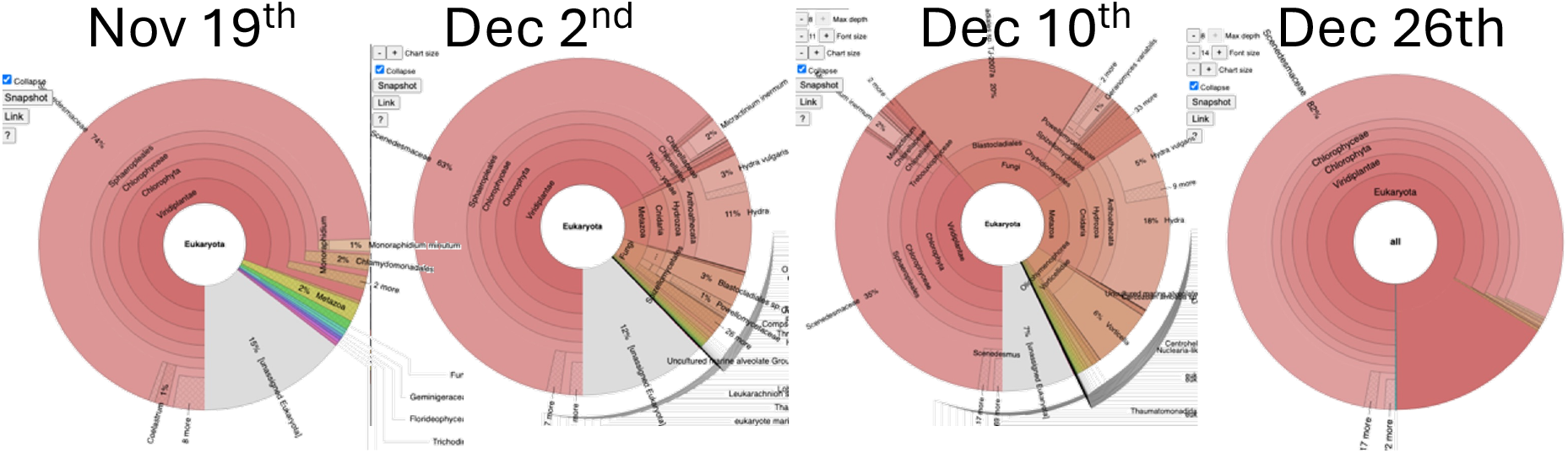
Krona charts of selected timelines from Pond 24 showing taxonomic classification of individual samples and the abundance distribution of contaminants. It shows that in the time line as of Nov 19th the majority of the reads were coming from the UTEX393. By December 2^nd^ we detect fungal and grazer contaminants.

Figure 5A shows a heatmap for abundances for prominent contaminant reads at the genus level during the 26B-AM SOT run during the same time period. We chose three ponds (represented as B9, B11, and B13, respectively), to do a detailed examination of the crash event. These contaminants noted in figure 4 are present in trace amounts only (<1% of total sequencing reads). While it is possible that biases in the extraction of genetic material are responsible for these low levels, it is also probable that fungi are more easily identified using the ITS gene rather than 18S. Major categories of contaminants in these ponds were *Fungi, Metazoa, Aphelida*, and other phototrophic contaminants. In Figure 5B, we show how these abundances compare in broad categories. These cultivation timelines were previously reported to have trace amounts of contaminants from the beginning of the timelines reported. Some of the samples were treated with the pesticide Fluazinam early on to remove pests. This allowed the ponds to recover from any infection. Following the timeline on the heatmap, it can be observed that in early November, we saw a higher relative abundance of Eukaryotic contaminants (although again represented by small read counts with <1% of total reads) than mid to late November for each of the ponds (B9, B11, and B13).

**Figure 5:**
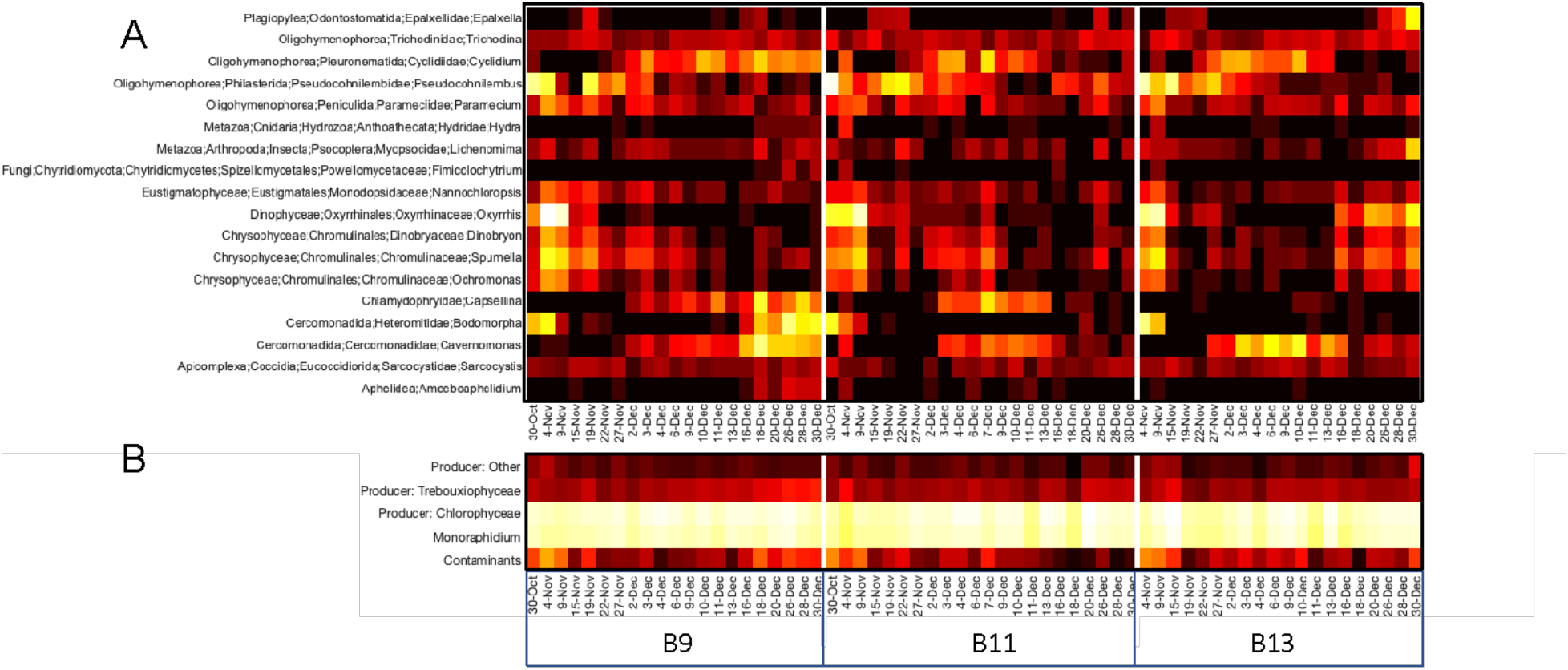
Classification of contaminant reads from the SOT cultivation runs of Monoraphidium minutum 26B-AM. We only get trace-amount of eukaryotic contaminants (<1% of total sequencing reads) A: shows a heatmap of log-scale abundances of reads at genus level of the contaminants, and B: shows a heatmap of log-scale abundances of reads for general broader categories.

Figure 6 shows Krona charts for the taxonomic abundance and distribution for a sample from December 30th to show that most of the reads were coming from the 26B-AM strain. We detected very low levels of contamination from grazers, flagellates such as *Cercomonadida*, and Aphelids. However, these pests are still present in trace amounts compared to algal 26B-AM read counts. This also demonstrate that detection of aphelids and other pests often implicated in crashes does not always lead to a crash occurrence [22].

**Figure 6:**
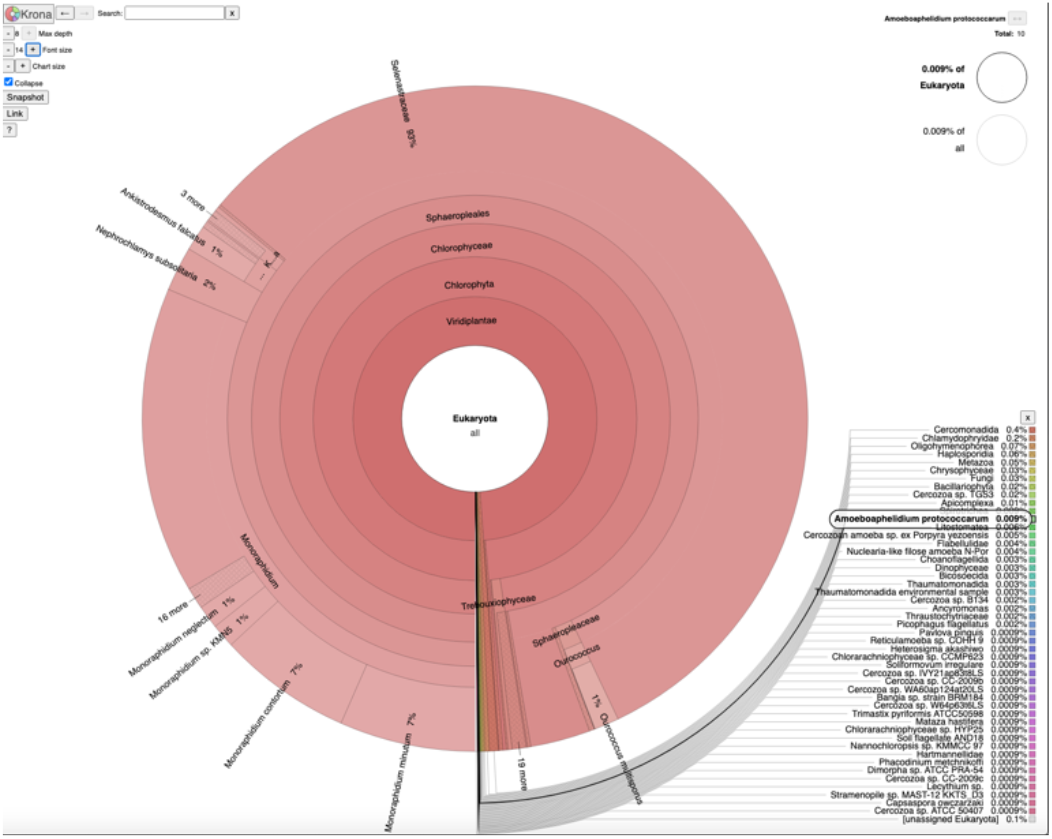
Krona chart for the distribution of the metagenomics reads for SPW9 ponds on Dec 30th showing the distribution of read abundance. 99% of reads are classified as algae species, and 93% of reads from the family containing Monoraphidium.

### Anomaly Detection Algorithm Update with Tensor Decomposition and Application on SOT Cultivation Data

Here we wanted to use the taxonomic abundance information to apply data science techniques to predict an oncoming crash in the absence of microscopy data. We have previously showcased a method for detecting an anomaly based on moving average computed on the ratio of sequence reads of contaminants vs. producers. We have upgraded this strategy to use higher-order multivariate joint statistics, specifically co-kurtosis, and its tensor decomposition to compute the measure of Hellinger distance as a measure of an anomaly[23]. In this method, we used the sequencing reads for producers and contaminants as an input for the algorithm, which decomposes data into several spatial sub-domains and timesteps. We compare the changes in these different components and their orientation and distance in this projected mathematical space. The Hellinger distance measures the orientation changes when new timepoints are introduced. This Hellinger distance was used as an anomaly score, and any significant changes in the score (>0.2 for SOT datasets) triggered the algorithm to call for an anomaly.

We successfully applied this method to the two available timelines of SOT cultivation runs, achieving early detection with > 3 days of early warning. This method predicted a crash for Pond 23 with a sufficient margin of early detection, which shows a 50% increase in the accuracy for detecting anomalies from the SOT cultivation runs. This shows successful application to the available SOT cultivation dataset, giving >3 days of early warning for crash. We also identified the major contaminants present in the ponds during this time period as chytrids, aphelids, and a metazoan hydra.

We used the sequencing reads for algae and contaminants as an input for the algorithm, which decomposes data into several spatial sub-domains and timesteps. We compared the changes in these different components and their orientation and distance in this projected mathematical space. The Hellinger distance measures the orientation and was recomputed when new timepoints were introduced. This Hellinger distance was used as an anomaly score, and any significant changes in the score triggered the algorithm to call for an anomaly.

We successfully demonstrated this method’s application to the three UTEX393 SOT cultivation runs, achieving early detection with more than 72 hours of early warning. We also applied this strategy for a 26B-AM strain cultivation run as a negative control, since this run had some contamination issues but recovered due to treatment. Figure 7 subplots A, B, and C show read counts of producer and contaminants over the cultivation timeline for the ponds that crashed and D, E, and F for uncrashed runs. This algorithm does not detect any anomalies in D, E, and F, which serve as negative control cases.

**Figure 7:**
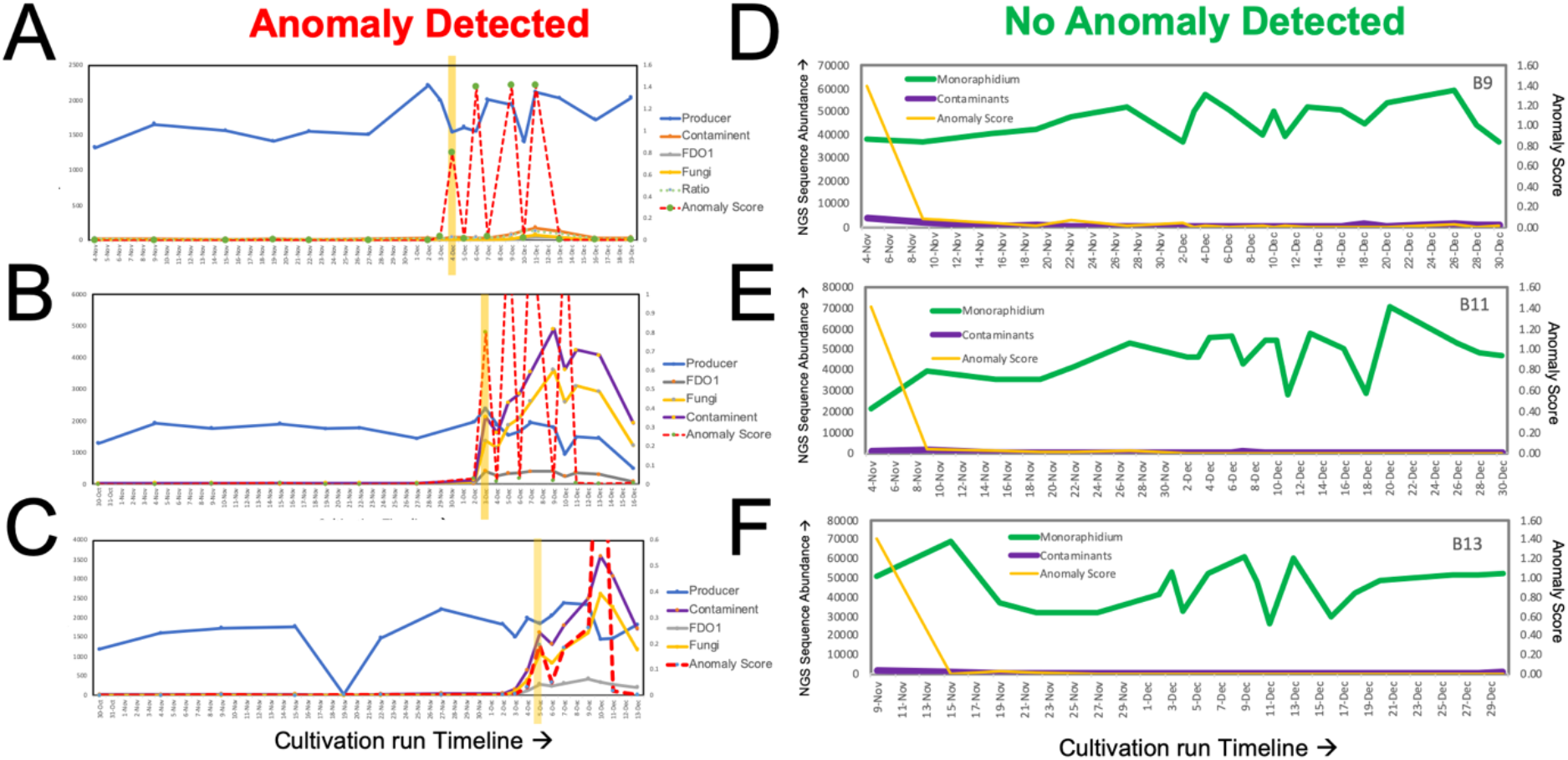
Anomaly detection algorithm PondSentry application on SOT cultivation NGS genus taxonomic abundance. A: Pond-23, B: Pond 24 and C: Pond 26 growing UTEX393, which had a pond-crash early Dec and D: B9, E: B11, F: B13, which did not crash. The time-series of raw counts of NGS reads of producers, contaminants, and computed anomaly score are presented in each subplot. This analysis showing PondSentry’s successful call for anomalies (crash) for time-series for crashed UTEX393 cultivation run and successfully not call anomaly in D, E, and F.

For timely detection of an impending pond crash we applied a novel anomaly detection method [23]. The method was originally proposed for distributed scientific computing data with solutions variables varying in space and time, but it is applicable to any setting involving multi-variate data sets with non-Gaussian distributions. The method does not label an individual sample as anomalous; rather it seeks to detect if/when some outlier samples are observed. It relies on the concept that for multi-variate data sets the information of anomalies is present in the higher-order joint moments and cumulants. Specifically, “principal kurtosis vectors”, computed by factorizing the co-kurtosis tensor (4th-order cumulant), identify directions in a multi-dimensional state space along which outliers are present. For a dataset of *N*_*v*_ variables 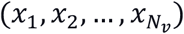 that are zero-mean (i.e. the mean subtracted from each), the co-kurtosis tensor of size *N*_*v*_×*N*_*v*_×*N*_*v*_×*N*_*v*_ can be expressed in index notation as

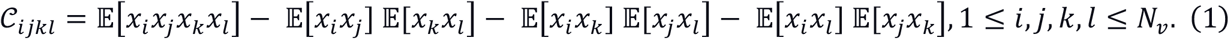

Here 𝔼 is the expectation (averaging) operation, and 𝒞 is a symmetric order-4 tensor (i.e. its entries are same for permuted indices). Accordingly, the principal kurtosis vectors are obtained by unfolding 𝒞 into a matrix of size 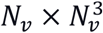 and performing a singular value decomposition.

This is analogous to the concept that the dimensions identified by principal component analysis (PCA), effectively eigenvectors of the covariance matrix, denote directions of maximal spread (variance). Accordingly, when even a few anomalous samples initially get added to an otherwise normal data set, it will result in a change in the orientation of principal kurtosis vectors (PKVs). Continuously computing and tracking the PKVs in appropriate space-time windows helps detect those in which PKVs suddenly change, indicating occurrence of anomalous samples in the corresponding window(s).

For the present application, we use taxonomic abundance information, i.e. amplicon sequencing reads of producers and contaminants, as the observational variables. Since each sequence represents a time snapshot of the pond, the data is purely time-varying with no spatial variation. A spatial variation could be analyzed if pond samples from multiple spatial locations were collected at each time and sequenced. This would allow identifying the spatial region of the pond where the crash originates.

For each pond we start with an initial time window of a certain size. The co-kurtosis tensor, and its PKVs, are computed using the data samples from that time window. For each successive temporal sample, the time window is expanded by adding the latest sample, and the co-kurtosis tensor and PKVs are recalculated. The new PKVs are compared against the previous PKVs, and when they change it indicates that the most recent sample contains anomalous sequences. Comparing one set of PKV to another requires a suitable measure; the method employs a “feature moment metric” (FMM), effectively the fraction of the overall moment (in this case the kurtosis) that is explained by each feature/variable. The FMMs are convenient since they sum to unity, and hence can be treated as discrete distributions, and compared using suitable divergence metrics, in this case a Hellinger distance. In summary each set of PKVs are translated to a corresponding set of FMMs, and computing the Hellinger distance between one FMMs set and another is equivalent to comparing the orientation of the PKVs.

### Feature Importance

We expanded the PondSentry method’s capability to multiple feature anomaly detection. Initial PondSentry predictions of the anomaly were based on simplified taxonomy time-series (producers and contaminants). For streamlining the detection of features triggering the anomalous event, we need to use higher resolution taxonomy features and identify the rank order of deleterious features responsible for the anomaly.

For this purpose, we used the time series of the abundance of three major players in the pond ecology for anomaly detection in UTEX393 strain cultivation (*Producer, Hydra, Fungi*). The feature “fungi” contain abundance reads aggregated for all fungal taxonomy combined. We are still using simplified taxonomy as we can’t expand the PondSentry method to more than five features due to the low number of time points in the time series. For the negative control, we used the 26B-AM SOT run in 2019, where the feature list used to apply PondSentry had five features (*Monoraphidium(26B-AM), Contaminants, Producer-Chlorophyceae, Producer-Trebouxiophyceae, and Producer-Other-Streptophyta*). We resorted to 5 features as the clustering of the reads can be explained by taxonomy.

The higher resolution of the PondSentry with multiple features was used for eukaryotic anomaly detection in raceway ponds and improved the early warning for detection of an anomaly by multiple days (greater than 72 hours). Also, with the measure of the principal kurtosis vector, we were able to determine the rank order of features responsible for the call of anomaly. We found that the feature “hydra” had the most contribution closely followed by “fungi” in the call for the anomaly in unialgal UTEX393 cultivation ponds.

### Prokaryotic signature of Pond Crashes for summer 2020 SOT cultivation runs

Samples taken between February and June 2020 were analyzed using 16S amplicon sequencing to obtain prokaryotic profiles for the pond samples. We utilized this dataset to obtain a general microbial profile for healthy ponds for top-performing DISCOVR strains. We used statistical methods to compare the microbial profile of specific pond samples collected from crashes (SPW12, May 15^th^ 2020, and SPW14, May 14^th^ 2020) with healthy samples. These pond crashes are due to bacterial contaminants, as explained briefly in this paper.

Figure 8 displays a general microbial profile of the ponds growing UTEX393 and 26B-AM, representing the top 95% of the sequencing reads in a heatmap with clustering columns (samples having a similar microbial profile to cluster together) and rows (taxonomy having correlated timelines close together). The samples’ microbial profile is correlated with the date of collection and the algal strain from our analysis. Secondly, May, June, and August’s microbial profiles were relatively steady for all the studied raceways.

**Figure 8:**
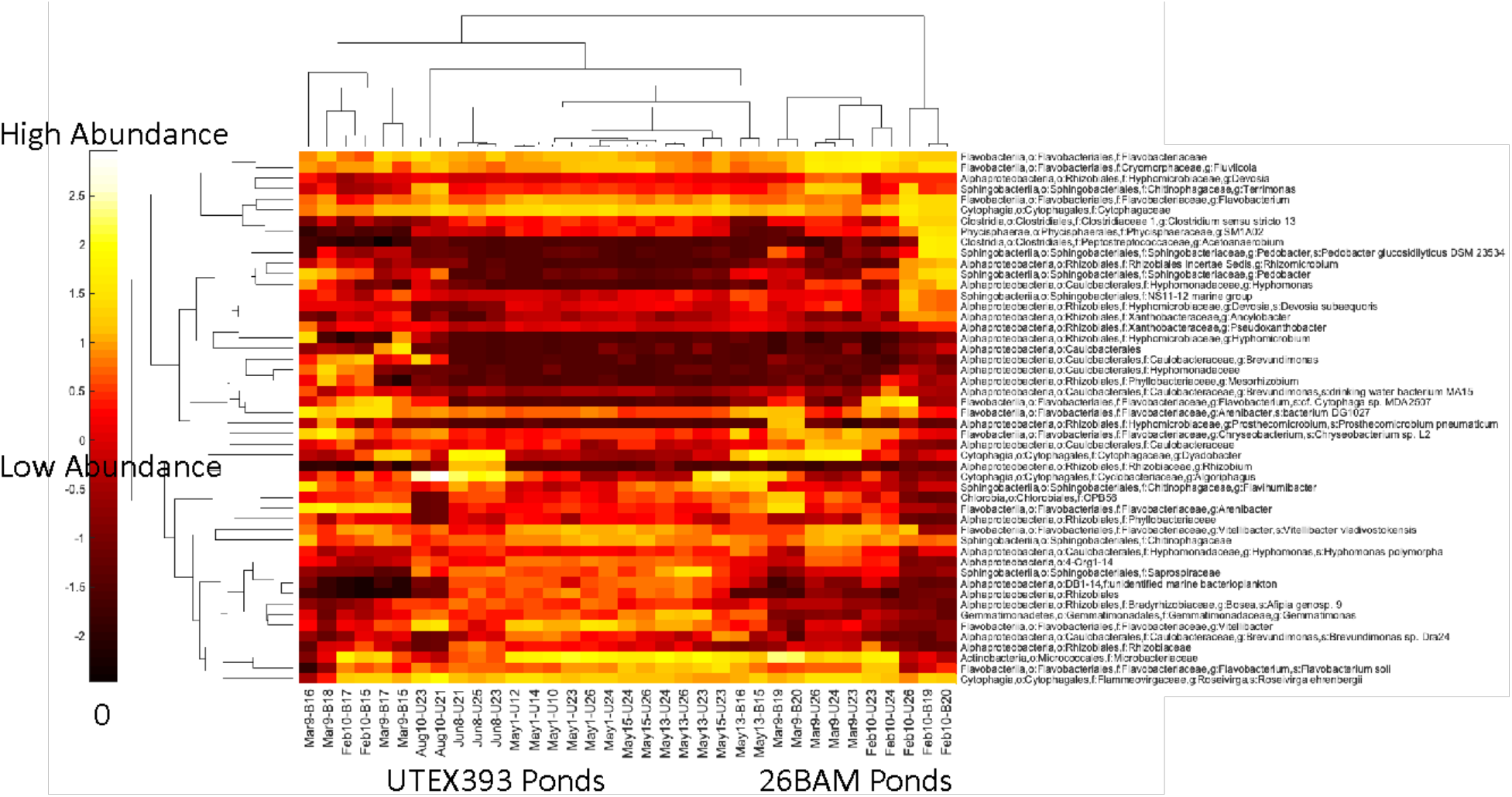
prokaryotic profile of ponds in AzCATI growing UTEX393 and 26B-AM representing top 95% of the sequencing reads.

In Table 2, we report the most abundant OTUs and their classification in the dataset. It is interesting to note that *Oligoflexus* and *Vitellibacter* account for almost 40% of the total reads in UTEX393 ponds, since *Oligoflexus* has been implicated in other raceway pond crashes. There is more diversity in the 26B-AM samples, with five OTUs making up for 50% reads.

**Table 1:** Estimated improvement in method with three feature anomaly detection.

| Method | Pond-23 | Pond-24 | Pond-26 |
| --- | --- | --- | --- |
| 2-feature anomaly prediction | 4-Dec | 3-Dec | 5-Dec |
| 3-feature anomaly prediction | 3-Dec | 27-Nov | 3-Dec |
| Estimated Improvement | +1 day | +6 days | +2 days |

**Table 2:**
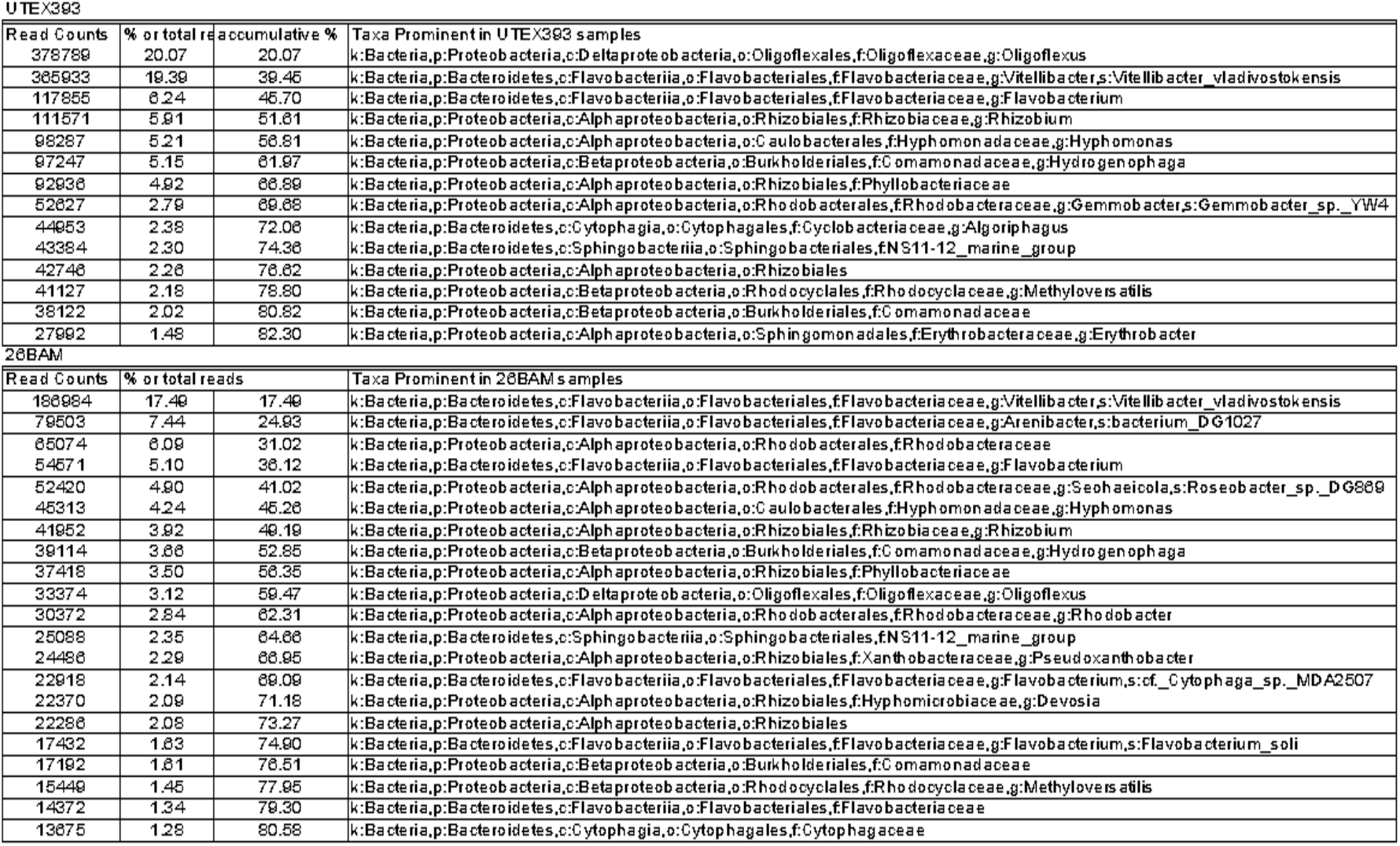
Table of Taxonomy for prominent OTUs which show >1% of the total read counts in respective ponds of UTEX393 or 26B-AM.

### Pairwise comparison analysis for identification of summer SOT prokaryotic crash signature

We used ponds samples which were reported to have a crash event (from SPW12, May 15^th^, 2020, and SPW14, May 14^th^, 2020) and, samples from healthy ponds (SPW10, May 13^th^, 2020, SPW10, May 15^th^, 2020, and SPW12, May 13^th^, 2020) for pairwise comparative analysis using the DESeq2 package in R. Figure 9 and Table 3 summarize the results from DESeq2 analysis.

**Table 3:**
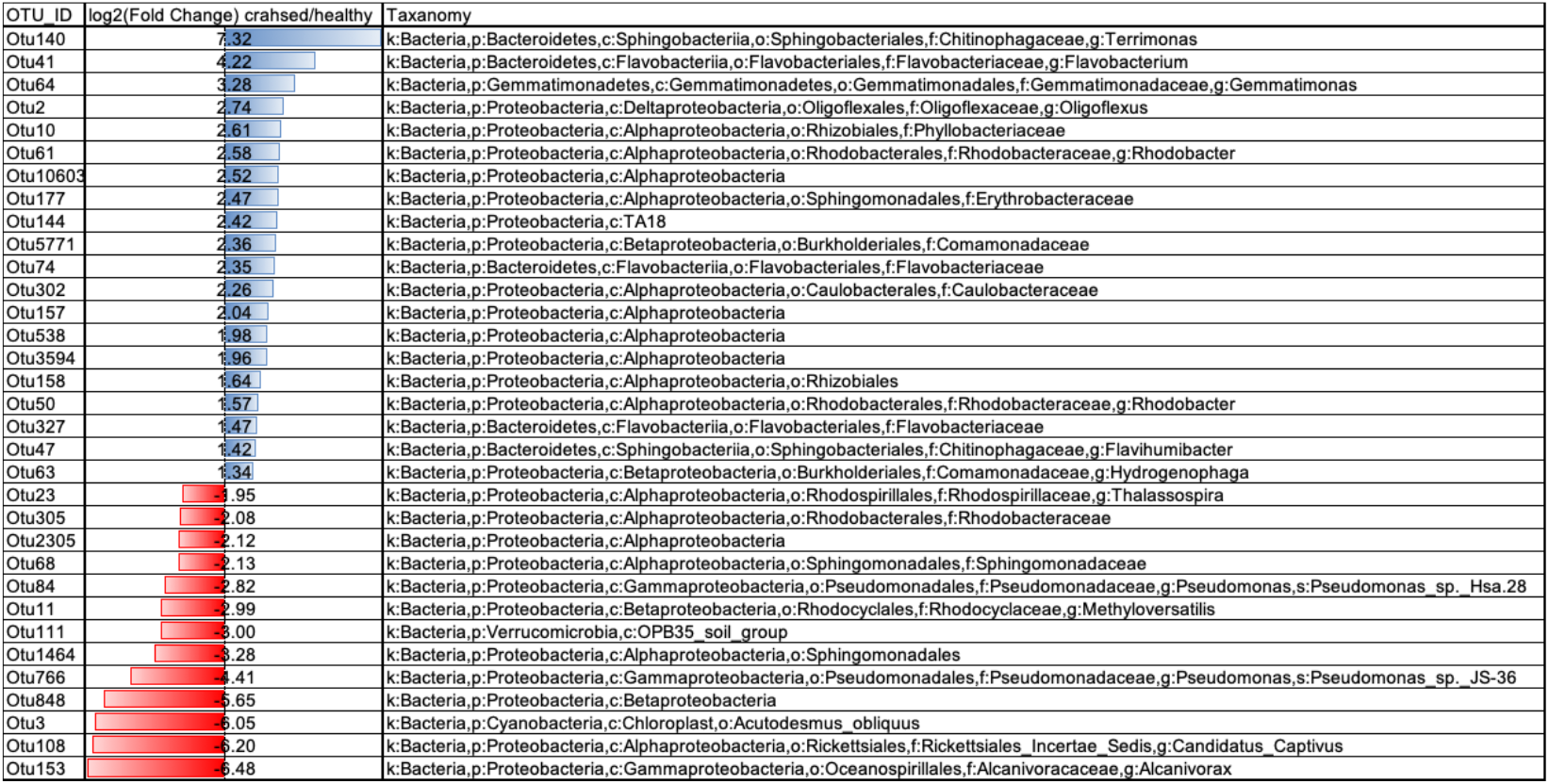
Table of OTUs and their classification, which shows significant changes in crashed pond samples from healthy samples using the DESeq2 algorithm.

**Figure 9:**
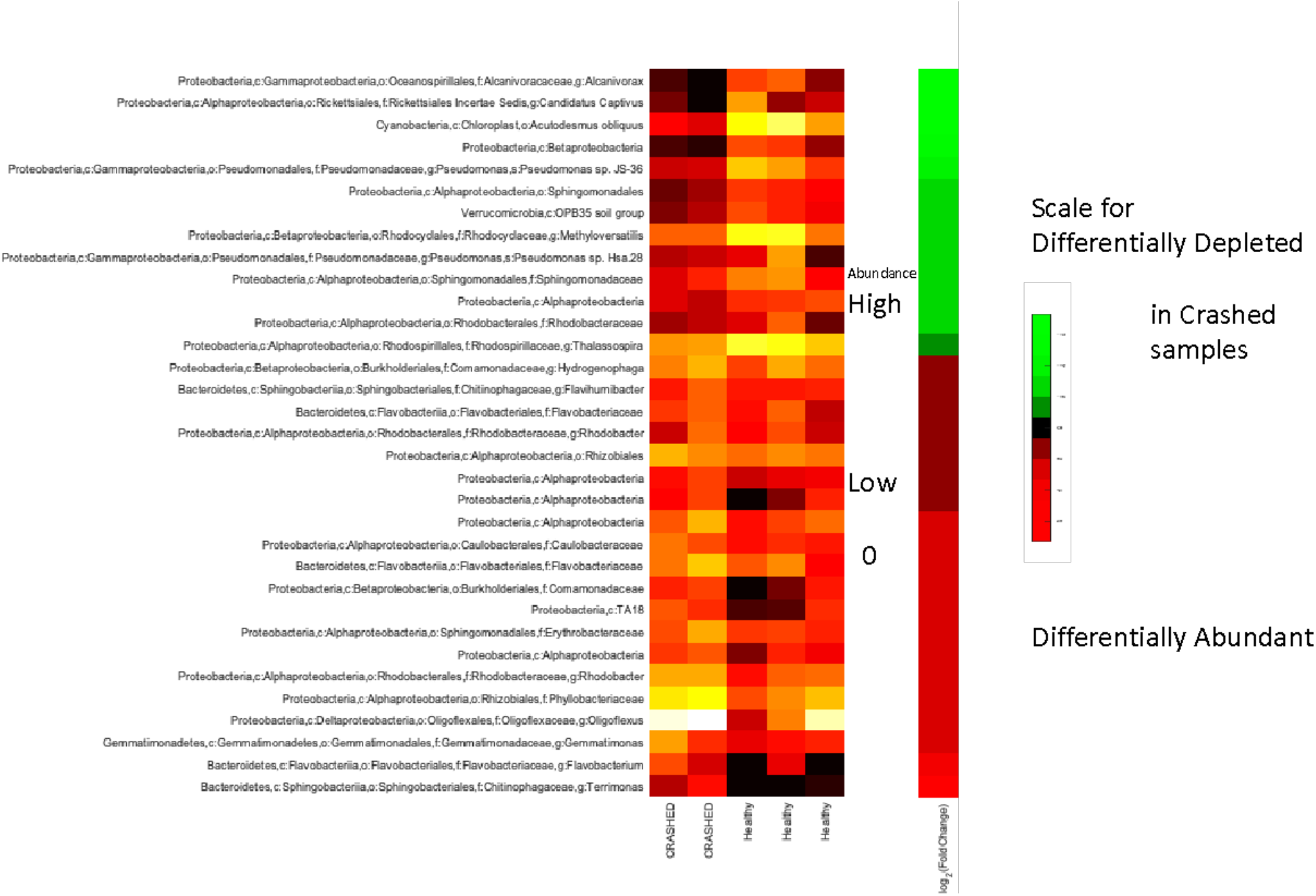
Heatmap displaying significant OTUs identified by Differential abundance analysis. The heatmap on the left shows the heatmap of abundance at each sample. The heatmap on the right shows the scale of change in abundance from crashed vs. healthy OTUs, with red being more abundant in crashed and green being more abundant in healthy.

Figure 9 displays read abundance for each sample on the heatmap on the left and the change in abundance (red-green scale) on the right. Green bands represent OTUs reads depleted in crashed samples and red being more abundant in crashed samples. The DESeq2 analysis computes significance in terms of p-values for the changes. We only present the significant taxa in Figure 9 and Table 3. We identify eight genera reported to be significantly higher abundance in the crashed samples, including the *Oligoflexus* strain responsible for the crash. While whole genome sequencing is not available for this data, *Oligoflexus* spp. have been reported as a causative agent for pond crashes during other cultivation periods[24].

## Conclusions

While microscopy and genetic methods can be used to get an overall picture of community composition in open air pond cultivation systems, initial infection by pests responsible for pond crashes is often at low levels difficult for humans to detect and below the detection level for qPCR probes. These probes are also blind to novel pests, which cannot be detected without whole-pond community composition sequencing. Amplicon sequencing of 16S and 18S genes allows researchers to observe both dominant and rare community members. However, determination of which of these members is a pest is much more difficult and requires time series data.

In the absence of a clear causative agent, our model, PestSentry, can give information on pests responsible for pond crashes, allowing researchers and industry partners to identify targets for pond protection methods. This method was used successfully to identify the community members responsible for a crash in a pond containing UTEX393 during winter cultivation trials at AzCATI, demonstrating clearly that a combination of chytrids, aphelids, and metazoans caused near-complete crop loss and lowered pond productivity. A non-crashed pond system was used as a negative control, confirming that although fungi were present in the non-crashed system, they were not identified as anomalous and are present at a low baseline level even in non-crashing systems. Without this method, microscopic identification of fungi would have led to treatment of the pond with pesticides, increasing the final biomass cost unnecessarily. Additionally, this method was used to determine the bacterial pest *Oligoflexus* as the causative agent for a different crash. This model can be used alongside traditional monitoring to limit pesticide treatment to times when it is most required to prevent crashes, leading to an overall lower biomass cost.

## Funding Acknowledgement

Sandia National Laboratories is a multimission laboratory managed and operated by National Technology & Engineering Solutions of Sandia, LLC, a wholly owned subsidiary of Honeywell International Inc., for the U.S. Department of Energy’s National Nuclear Security Administration under contract DE-NA0003525. This work was funded by the US Department of Energy’s Office of Critical Minerals and Energy Innovation under the DISCOVR project, contract NL0040168. This paper describes objective technical results and analysis. Any subjective views or opinions that might be expressed in the paper do not necessarily represent the views of the U.S. Department of Energy or the United States Government. This article has been authored by an employee of National Technology & Engineering Solutions of Sandia, LLC under Contract No. DE-NA0003525 with the U.S. Department of Energy (DOE). The employee owns all right, title and interest in and to the article and is solely responsible for its contents. The United States Government retains and the publisher, by accepting the article for publication, acknowledges that the United States Government retains a non-exclusive, paid-up, irrevocable, world-wide license to publish or reproduce the published form of this article or allow others to do so, for United States Government purposes. The DOE will provide public access to these results of federally sponsored research in accordance with the DOE Public Access Plan https://www.energy.gov/downloads/doe-public-access-plan.

